# Machine learning segmentation tool trained on synthetic data for tracking cytoskeleton polymerisation and depolymerisation

**DOI:** 10.1101/2025.01.04.631322

**Authors:** Karan Elangovan, Tamsin A. Spelman, Gilles Dupouy, Gaurav Singh, Marie-Edith Chaboute, Henrik Jonsson

## Abstract

The cytoskeleton is important in controlling the growth and morphology of plant cells, so tracking its morphological changes is essential. Here, we develop a new machine learning based segmentation tool for microtubules (MTs), which can distinguish between polymerised and depolymerised fibres. To circumvent the low abundance of data, we trained on synthetic images of microtubules from a computational microtubule model, pre-processed to reproduce microscope effects and partial depolymerisation. We used this tool to investigate how the MT network in an *Arabidopsis thaliana* root hair cell repolymerises after depolymerisation under Oryzalin (OZ) drug treatments. Specifically, we show the network initially repolymerises from the shank region. This work demonstrates the viability of using synthetic data to train machine learning systems handling cytoskeletal image data.

## 1 Introduction

The cytoskeleton is found in most cells and is fundamental to cellular function including organelle transport; structural framework and cell division [1]. The plant cell cytoskeleton consists of two fibres: microtubules (MTs) and actin. This is an active network with the fibres constantly growing, shrinking and interacting, allowing organised networks to form. The anisotropy of the microtubule network depends on microtubule interaction dynamics [2,3]. It also depends on cell shape, as experiments in protoplasts showed microtubules aligning along the long axis of the cell [4]. However, microtubules do not always align along the long axis of the cells, for example in growing Hypocotyl cells the microtubules align transversely along the cell [5]. It has been proposed that some microtubule reorganisation could be related to external cues such as mechanical stress or light [6]. Even at a more local level microtubules can form local bands [7]. Cortical microtubule behavior has been modelled with multiple frameworks including Tubulaton [8, 9], CorticalSim [10] and Cytosim [11]. However, to observe these different behaviours and organisation we need powerful segmentation tools to study the cytoskeleton network quantitatively.

Segmentation tools have been developed to segment plant cells in different plant tissues [12–14]. These adopt both traditional and more recent machine learning tools. Machine learning tools trained on synthetic data in particular have shown recent promise. Human retinal network have been extracted from optical coherence tomography angiographs using a generative adversarial network trained on synthetic networks generated by physics model [15]. Separately, SyMBac was developed to generate synthetic images of bacterial cell colonies for training machine learning algorithms (specifically DeLTA and Omnipose) to segment cells from experimental images of bacterial colonies [16]. Specifically algorithms have been developed to segment the cytoskeleton [17]. While many of these tools are not published open source, making these methods difficult to utilise with new data, some of them such as FilamentSensor2.0 [18], CytoSeg2.0 [19] for actin filaments, SOAX for biopolymer networks [20], JFilament implemented in Image-J [21] are made available. However, developing a universal tool is also challenging, meaning existing tools may not generalise to new data quality. As such, these algorithms can struggle in particular settings, for example detecting depolymerised microtubules (Fig 1).

**Figure 1:**
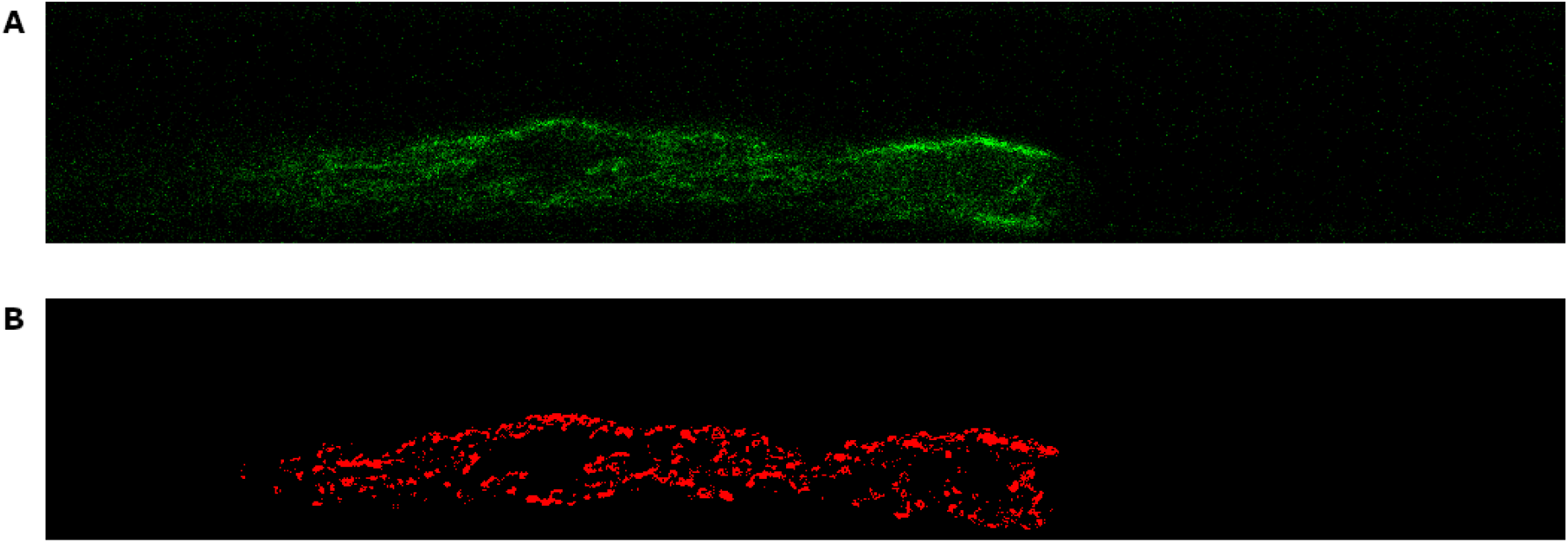
The results of segmenting a fully depolymerised image using a standard pre-existing technique (Ilastik) **(A)** The original image **(B)** The predicted segmentation

The arrangement of microtubules within the *Arabidopsis thaliana* plant root hair cell consists of a cortical network and endoplasmic microtubules [22] and there is evidence that microtubules nucleate off from the nucleus [23]. This changes markedly at maturity where the microtubule network becomes much more sparse [22]. There is also an active actin network in the cell [24]. The root hair itself is a long, thin tip growing cell which grows off the trichoblast cell on the edge of the root, reaching about 1mm in length and about 10*µ*m in diameter [25]. During growth the nucleus migrates up the root hair which is necessary for growth [26]. Using a microchannel setup [27] the relationship between nucleus at different stages of development have been explored, and the impact by cytoskeleton drug treatments [28] have demonstrated the cytoskeleton plays a key role in this behaviour. However, to understand how local changes in the cytoskeleton are effecting the nucleus and thus growth of this cell under depolymerisation conditions, a segmentation tool is needed to quantitatively detect cytoskeleton fibres and their depolymerisation. To our knowledge, no previous studies have looked at tracking these changes with difficulties arising as diffuse residue is still present and detectable making intensity threshold for detection challenging.

We develop a machine learning based tool trained on synthetic data to segment the microtubules, which can distinguish between polymerised fibres and depolyermised fibres (and background). We use this to track the depolymerization in plant root hairs. In Sec 2.1 we outline our pipeline for generating synthetic microtubule images to use for training data followed by the architecture of the machine learning algorithm in 2.2 and a brief outline of how experimental data was obtained in 2.3. In Sec 3 we discuss the outcomes from the model. We compare the synthetic data generation to experimental data in 3.1. We compare our method on polymerised and depolymerised data in Sec 3.2–3.3 and use it to track cytoskeleton recovery in Sec. 3.4. Finally we conclude in Sec 4.

## 2 Materials and methods

To track the microtubule depolymerisation we use a convolutional neural network following the standard U-Net [29] architecture (Fig 2). However, plant microscopy data are slow and expensive to collect, which limits the availability of training data. In particular, marked training data generally require experts to annotate. Therefore, we first develop a pipeline to generate synthetic training data (Fig 2).

**Figure 2:**
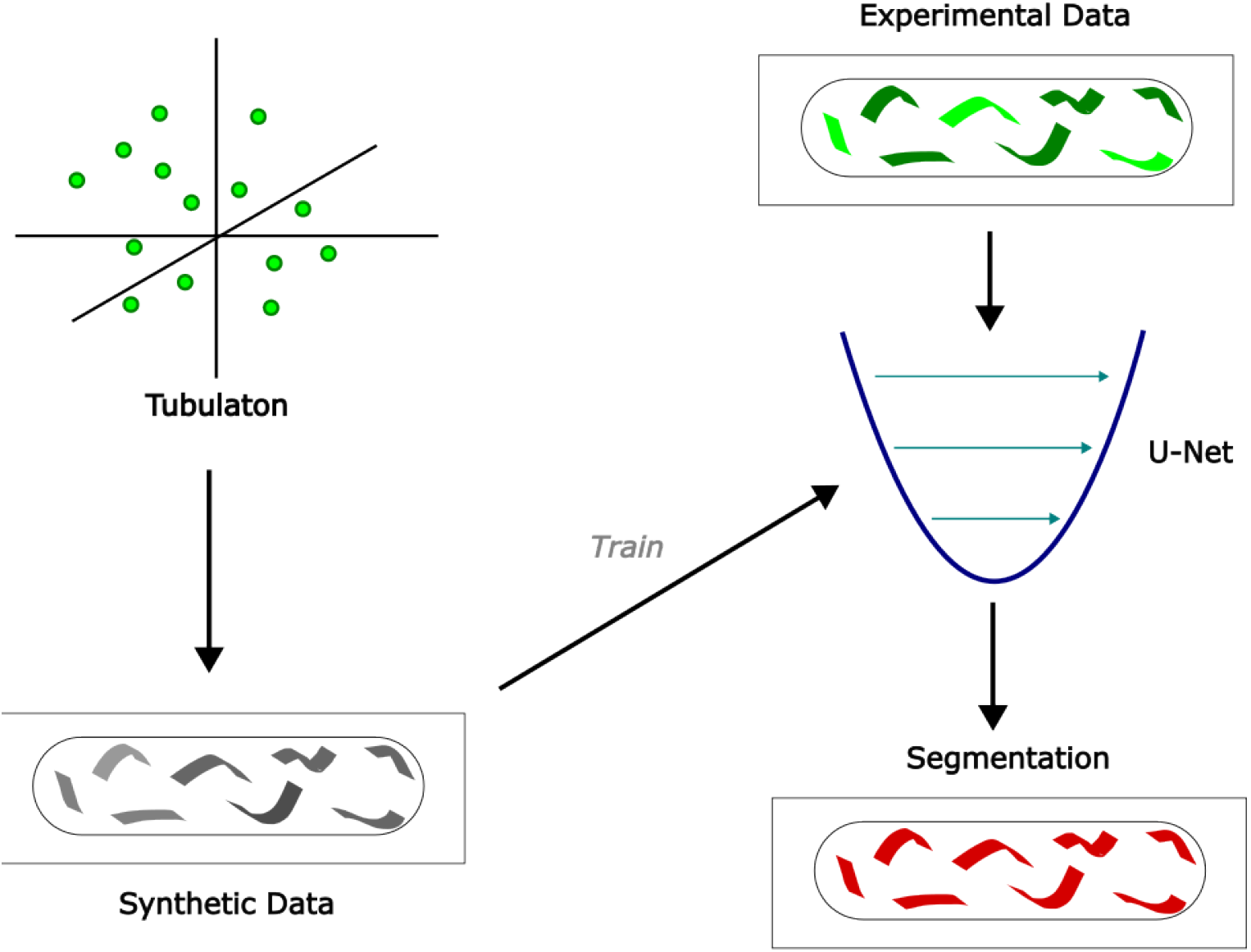
Outline of the synthetic data generation and segmentation pipeline.

### 2.1 Synthetic data generation

To generate the synthetic training data, we used existing MT modelling software to generate realistic MT networks where the exact position of every MT is known, then post-processed the images to imitate microscope images (Fig 3).

**Figure 3:**
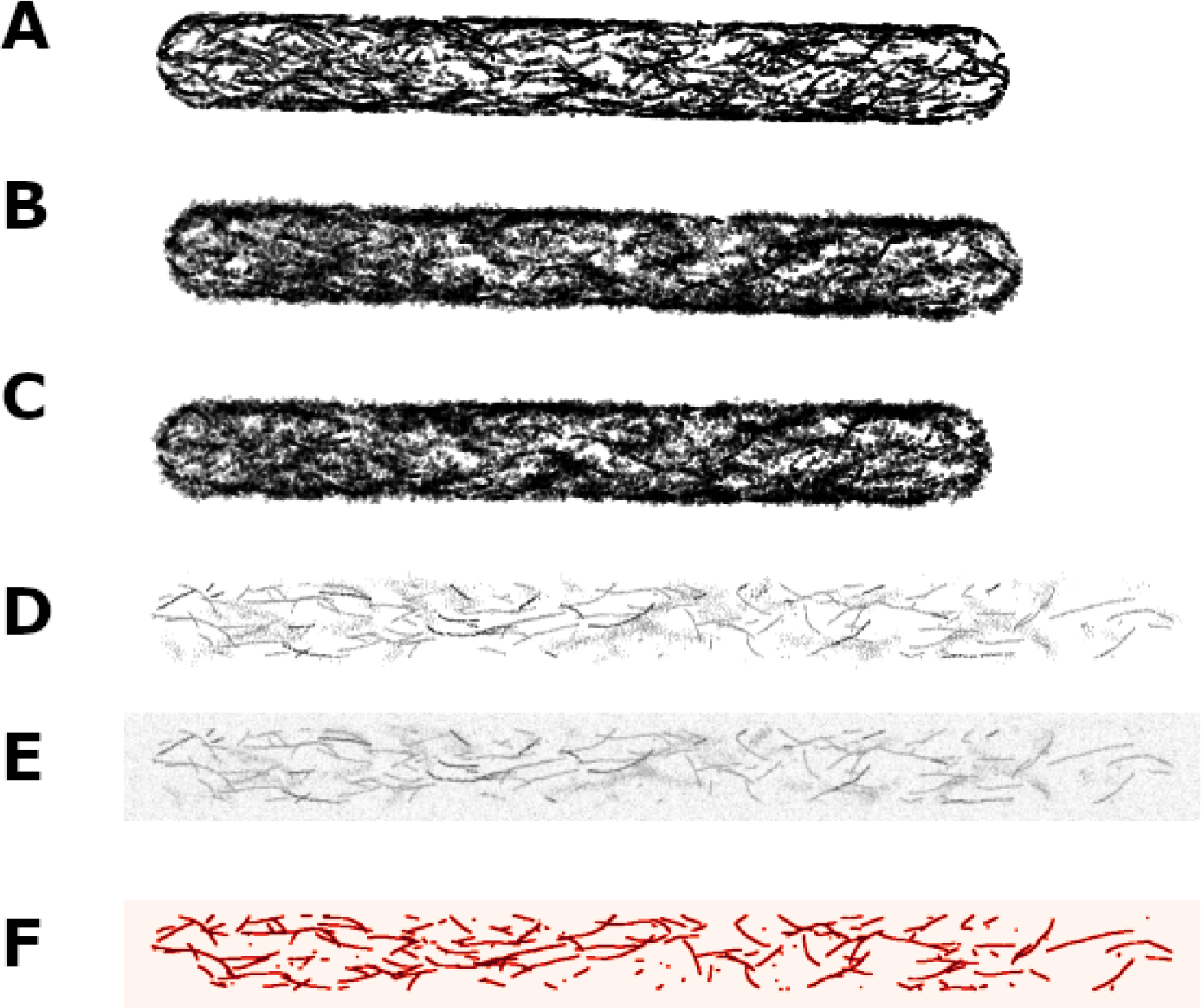
Examples of image at main stages of the synthetic data generation pipeline. **A**, The original Tubulaton output (3D) **B**, After simulating depolymerisation (3D) **C** Simulated positions of the FPs (3D) **D**, The projection of the FPs onto a plane **E** After convolving with point spread function (This is the final image) **F** The corresponding mask used as ground truth

**Figure 4:**
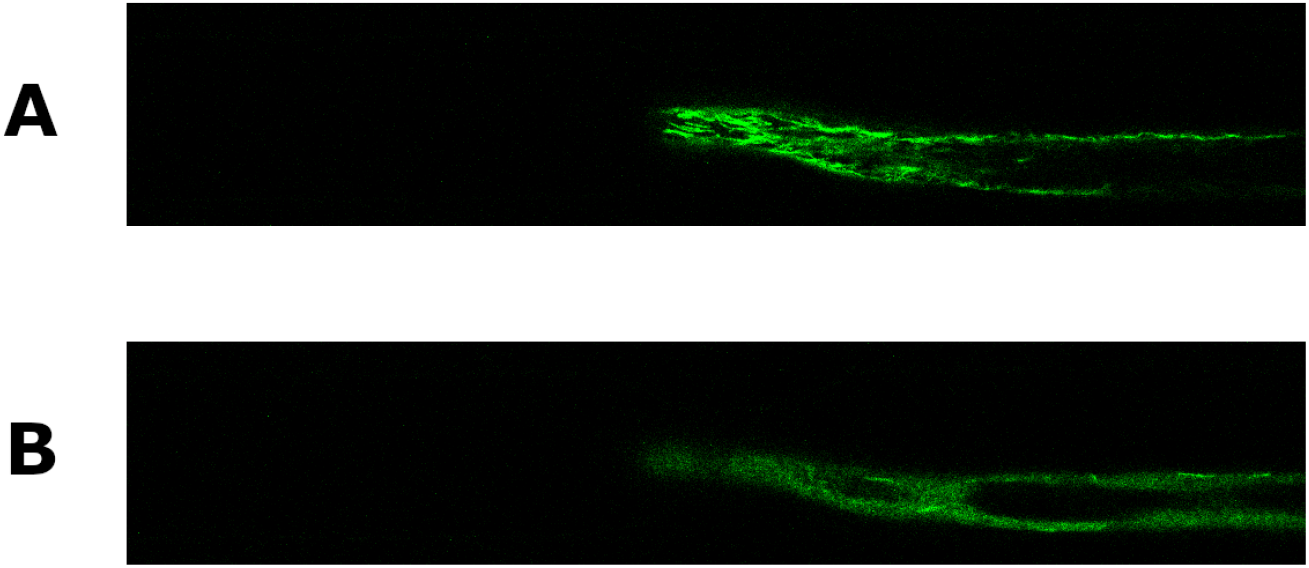
MT network, tagged with SUN2-MBD, in *Arabidopsis thaliana* root hair cell, with contrast adjusted. **(A)** Before oryzalin treatment **(B)** After oryzalin treatment.

The simulated microtubule networks were produced with Tubulaton [4, 8, 9]. In brief, Tubulaton models every microtubule in 3D separately as a series of unit vectors placed end to end, capturing growth, shrinking, and tread-milling by adding or removing unit vectors from the microtubule plus or minus end every time step (Fig 3A). Unimpeded microtubule growth direction is a normalised weighted summation of its current direction, an imposed bias direction and a small random component. Microtubule behaviours such as nucleation, spontaneous catastrophe and dis-attachment are incorporated. Additionally, microtubule interactions such as bundling, induced catastrophe and crossover severing are also modelled. The cell shape is set by a prescribed surface mesh. Microtubule growing ends which reach close enough to the cell surface, either undergo catastrophe at large interaction angles or at small interaction angles change their growth direction primarily along the cell surface (with a small random component) to prevent them leaving the cell.

To produce simulated microtubule networks similar to the long, thin root hair, we prescribe the cell shape as a cylinder with hemispherical capped ends. Microtubule nucleation occurs on the cortex of the cell only. We also apply a small bias to the microtubule direction, so they preferentially favour the long axis of the cell. Experimentally, microtubules preferentially align along the long axis of the cell [4]. In our simulations, the bias allows this feature to appear more quickly in the network after initiation from no microtubules, thus decreasing simulation time. This is sufficient for generating synthetic data as we need our networks to visually appear similar enough to the microscope images, but do not require them to reach dynamical equilibrium.

The Tubulaton output gives the exact location of every simulated microtubule, with each microtubule individually identifiable (Fig 3) However, upon projection this produces a binary image so post-processing is required to more closely match experimental images. The post-processing reproduces depolymerisation, fluorophore effects, and microscope effects.

To mimic depolymerised MTs in the synthetic image, we imitate dissociated monomers undergoing Brownian motion, by spreading out microtubule positions. Each Tubulaton outputted MT is depolymerised with a probability *p*_*d*_. To each point within selected MTs to depolymerise we add an isotropic Gaussian displacement. The probability of depolymerisation *p*_*d*_ is varied between images, with the *p*_*d*_ for each image varied to ensure diversity of synthetic training data (specifically, we used *p*_*d*_ values 0.2, 0.5 and 0.8) Because of the large number of fluorophores (FPs), modelling each individually would be too computationally expensive. So instead we group FPs into equal sized FP “packets”. These smaller number of packets are then assigned randomly without replacement to each MT unit vector (Fig 3C).

We next produce a 2D image by projecting all the FPs orthogonally, but only taking FPs in a specific z-range interval, as FPs far away from the focal plane would not show up in a confocal microscope image (Fig 3D). The centring of the interval is varied between different images in the same dataset.

We simulate further variation in brightness by scaling the pixel intensities of each MT uniformly, by a random amount (with the distribution being a tunable hyperparameter).

We mimic blurring caused by the confocal microscope via convolution with an Airy Disc kernel, which has formula:

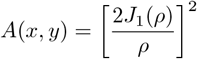

where 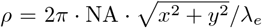, NA is numerical aperature, *λ*_*e*_ is emission wavelength of the FPs, and *J*_1_ is the Bessel function of the first kind of order 1 (Fig 3E).

To ensure that our synthetic images more closely match the lower resolution of the experimental images, we downsample. We break the image into 10 *×* 10 tiles and pad tiles at the edge of the image (i.e. on the right and bottom) with additional rows and columns of 0s to deal with overflow. Each tile is replaced with a single pixel of intensity taken at random for the tile, to produce a lower resolution image.

We create the ground-truth label for the synthetic image by referring back to the orthogonal projections of (some of) the FPs onto the focal plane. Then we downsample with the same tiling technique, this time taking the maximum of each tile instead of a random sample (i.e. we label a pixel as a polymerised MT if the corresponding tile has any FPs).

We perform one last noise operation to the image only, to account for any further residual noise effects. For each pixel, the new intensity is:

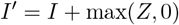

where *I* is the original intensity and *Z* ∼ Normal(0, *σ*^2^), and *σ*^2^ tunable.

Finally, we renormalise the intensities to make the maximum value 1.

This process has transformed the binary output of Tubulaton, where each pixel value represents either the presence or absence of a microtubule, into an image with a continuous spectrum pixel intensities, more closely resembling an experimentally obtained microtubule image.

### 2.2 Segmentation pipeline

Now we can generate large quantities of synthetic training data, we use it to train a machine learning tool to segment the microtubule images to detect polymerised microtubules. The model will produce a probability that each pixel of the image is part of polymerised microtubule. This can then be converted to a binary mask by setting a hard threshold above which we assume the pixel corresponds to a polymerised microtubule (Fig 3F). We use the threshold of 0.5.

The large size of the experimental images and equivalent synthetic images, made training with these whole images very slow. Exact pixel dimensions also varied between experimental images, so the segmentation method needed to be flexible enough to cope with this variability. Therefore, rather than training on the whole image, we trained using square patches of 128 × 128 pixels extracted from the synthetic data.

After training the model though, when running the segmentation on experimental images, we ran the segmentation on individual patches but needed to be able to recombine patches to produce a segmentation which corresponded to the original experimental image. However, using regular non-overlapping patches produces artifacts at the boundaries of the arbitrarily chosen patch boundaries. Therefore, for inference the model is run on chequerboard offset patches that wrap around the right and bottom of the image, so each pixel is in two patches. We take he final predicted probability that each pixel represents a microtubule as the average of the two values produced for that pixel.

We used a convolutional neural network for the semantic segmentation, specifically a U-Net architecture [29]. For training we used 5800 synthetic patches generated from 580 separate synthetic images. We trained the model for 500 epochs using an Adam [30] optimizer, with a batch size of 3, drop out rate of 0.1 and a learning rate of 3 *×* 10^−6^. We used weighted binary cross entropy loss with a 9 : 1 weighting scheme, to account for the severe class imbalance between polymerised MT pixels and background.

The network was implemented in PyTorch in Python 3.11.8. The code used to run the simulations discussed in this paper is available at the Sainsbury Laboratory Software repository https://gitlab.developers.cam.ac.uk/slcu/teamhj/publications/elangovan_etal_2025

### 2.3 Experimental images of root hairs

Images of the polymerised and depolymerised microtubules *Arabidopsis thaliana* root hairs used for testing our ML algorithm, were collected using the microchannel setup outlined in [27]. *Arabidopsis thaliana* Col-0 seedlings expressing p35S:GFP-MBD microtubule marker (and a pSUN2:SUN2-tagRFP nucleus marker) were observed under a LSM700 confocal microscope. To view microtubule depolymerisation the liquid media within the microfluidic chip was replaced with 1*µ*M of Oryzalin which depolymerised the microtubule network.

## 3 Results

### 3.1 Comparison of synthetic data to experimental data

A synthetic dataset used for training the neural network was generated using a pipeline combining several steps (Fig 3).

The synthetically generated images capture features of the experimental images (Fig 5). In fully polymerised data most of the MTs are clearly visible as solid colour although some sections with a different texture in the experimental images suggest some fluorescence is still visible (Fig 5A). This is reflected in the synthetic data. In experiments microtubules can be depolymerised using Oryzalin treatment, which can be captured in the synthetic data by increasing the depolymerisation probability *p*_*d*_. This results in less well-defined microtubules and a spread of fluorescence caused by the depolymerisation (Fig 5B). After 10 minutes in experiments, the microtubules have fully depolymerised and we observe a more spread out fluorescence (Fig 5C). This is similarly reflected in the synthetic data caused by using a *p*_*d*_ value close to 1. However, there are visible differences between the synthetic and experimental data (Fig. 5). In the experimental data we observe some dark regions with no fluorescence, which are likely related to the presence of organelles such as the nucleus or vacuole which prevent MTs from entering. For simplicity, this is not simulated in the synthetic data generation.

**Figure 5:**
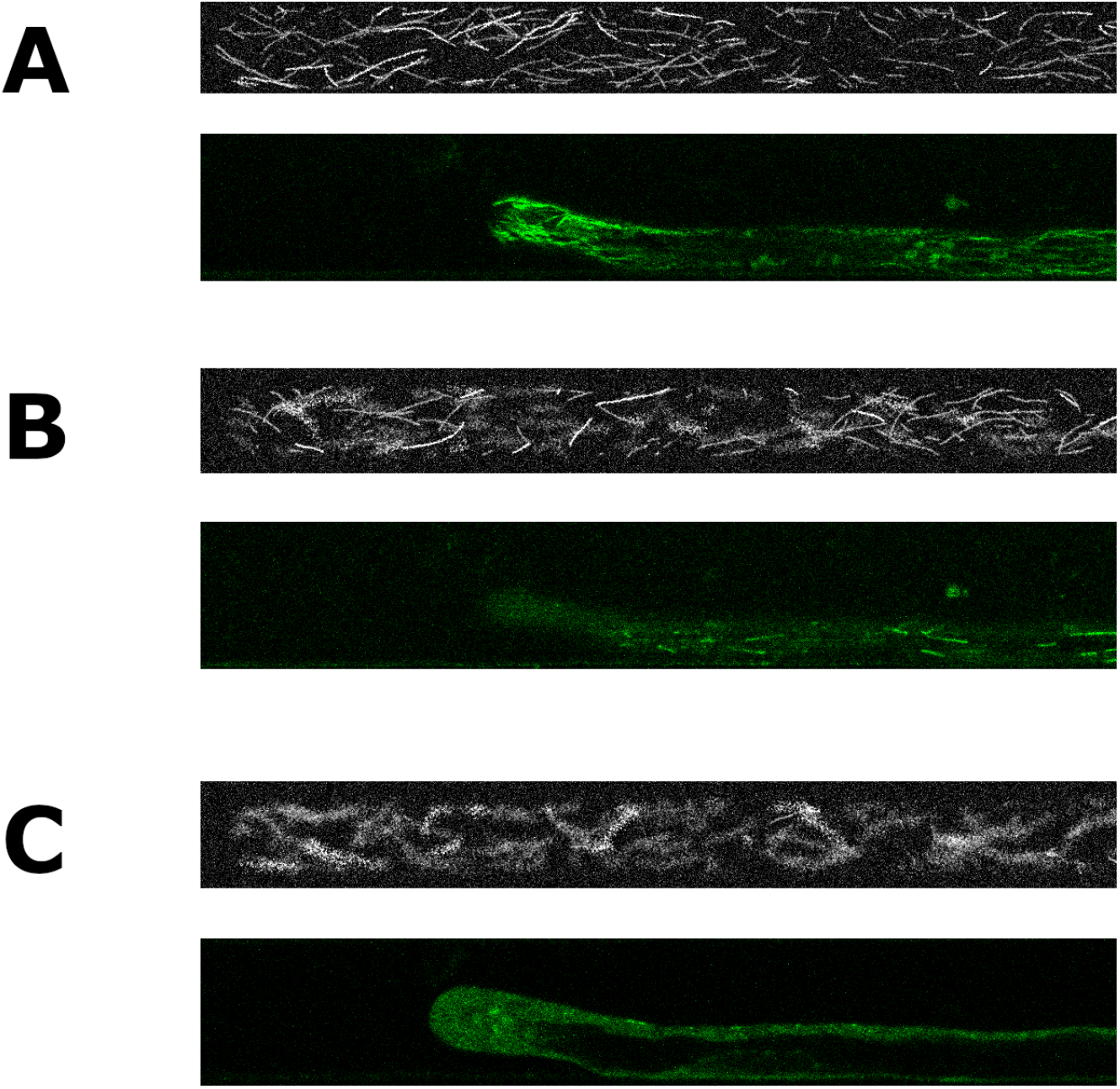
Representative examples of synthetic data compared to experimental data for three cases **A**, No depolymerisation **B**, “Partial” depolymerisation (*p*_*d*_ = 0.5 for synthetic) **C**, Total depolymerisation. In each subfigure, the top panel displays example synthetic data and the bottom panel the experimental image (contrast adjusted for visibility).

### 3.2 Outcome from segmentation

To obtain an indication of the ability to segment data of various degrees of polymerisation, we analysed representative segmentations from the model on specific images, covering the case of a dense polymerised network, a sparsely polymerised network, and a depolymerised network (Fig 6).

**Figure 6:**
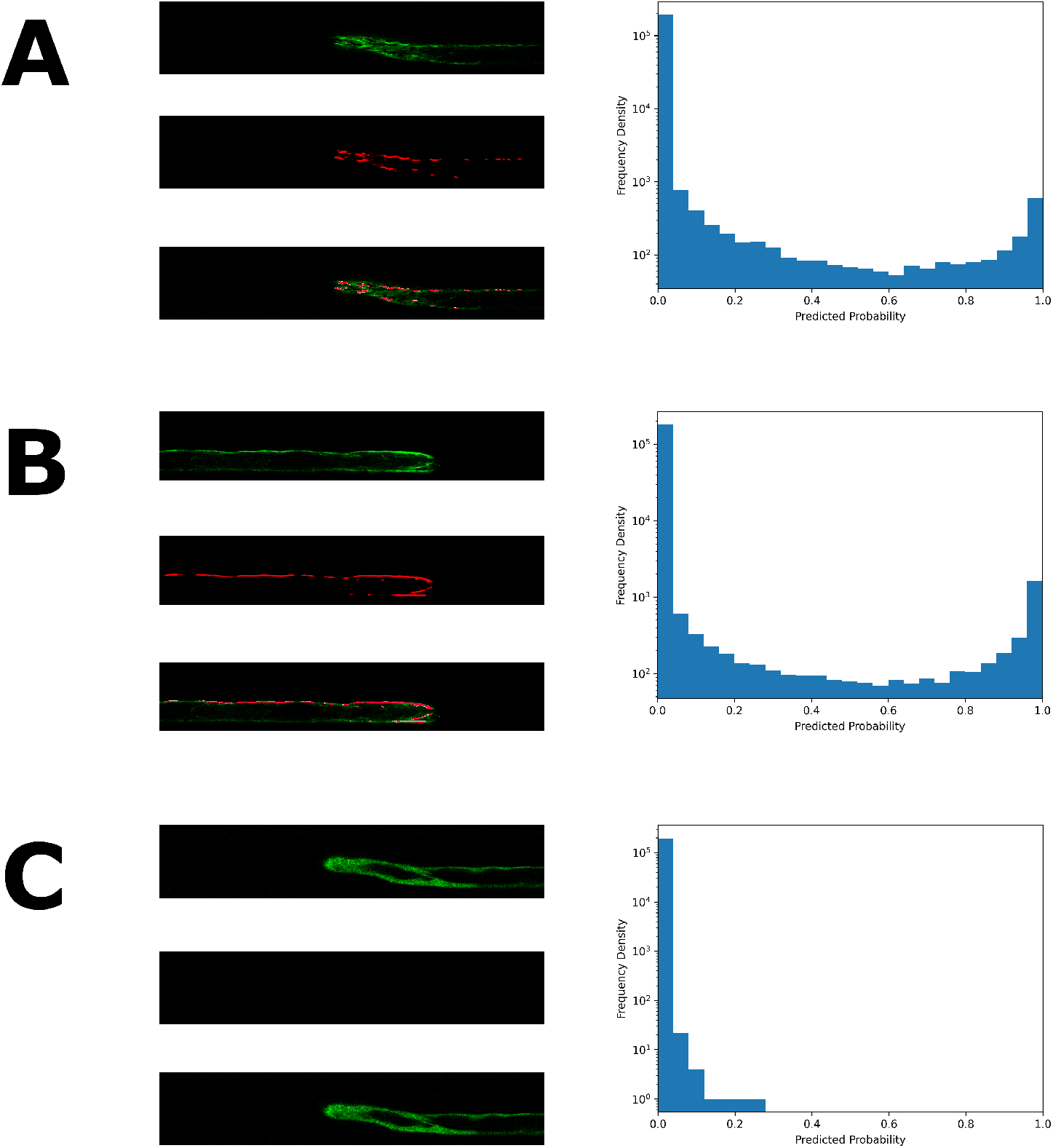
The trained model’s segmentations of experimental images for a **(A)** dense network **(B)** sparse network **(C)** depolymerised network. In each subfigure the lefthand images from top to bottom show: the original experimental image, the soft segmentation predicted by the model, the predicted hard segmentation (red) overlaid the original image (green). The histograms show the distribution of pixel predicted probability of being part of a polymerised MT.

The “soft segmentations” are given by taking the sigmoid of the model’s outputs (the model predicts logits instead of probabilities), and give a value between 0 and 1 to each pixel representing the probability that pixel is part of a microtubule. Here 0 or 1 representing absolute certainty the pixel is background, or polymerised MT, respectively. The “hard segmentation” is a binary image showing the pixels predicted to be polymerised and is decided from the soft segmentation by using a threshold of ≥ 0.5 for a pixel to be polymerised.

As we see in the histograms, almost all pixels are assigned probabilities close to 0 or 1 (note the log scale), showing the model is very sure in most cases, and so robust to minor changes to the arbitrarily chosen threshold of 0.5.

### 3.3 Evaluation of segmentation on polymerised and depolymerised data

To evaluate the effectiveness of our model trained on synthetic data we test it on two data sets representing completely polymerised and completely depolymerised MTs in root hair cells. The first set is of untreated experimental control images. We generate a ground truth by segmenting using Ilastik [31]. Using Ilastik involves significant human involvement to mark some data manually and re-check images at each step when the algorithm updates. In the fully polymerised regime, the positioning of MTs is clearer, so we use a segmentation generated with Ilastik against which to compare our model. Note that this pipeline optimised for control data performs poorly on depolymerised data (Fig. 1). The second set of experimental images are taken 10 minutes after treatment with Oryzalin, so we expect the microtubule network to be fully depolymerised. Thus, we take the ground truth as the whole image being background, i.e. no microtubules present. We evaluate these against the hard segmentation taken from its probability predictions on each pixel by thresholding at probability 0.5.

We validate using a variety of standard metrics for binary classification. The results for the control data are shown in Table 1 and for the depolymerised data in Table 2. We cannot validate on partially depolymerised experimental data as we have no reliable technique for getting ground truth segmentations in this case, we assume that the model generalises correctly to such images as it is accurate at both extremes (unlike classical techniques that do not work in the totally depolymerised regime) For the control data we obtain a good value for the AUC (area under curve) for PR (precision-recall) and ROC (receiver operating characteristic). There is a class imbalance of about 1:9, so the scores achieved are good, relative to the benchmark of 0.1 by just predicting each pixel with the the mean probability.

**Table 1:**
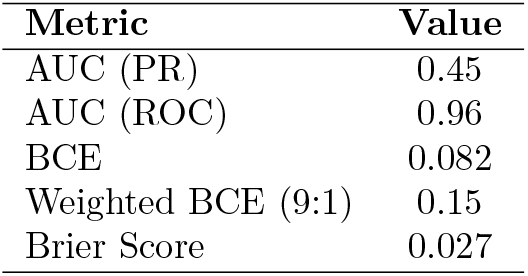
Performance of the model on an experimental dataset of *n* = 148 control images, using Ilastik segmentations as ground truth.

| **Metric** | **Value** |
| --- | --- |
| AUC (PR) | 0.45 |
| AUC (ROC) | 0.96 |
| BCE | 0.082 |
| Weighted BCE (9:1) | 0.15 |
| Brier Score | 0.027 |

**Table 2:** Performance of the model on an experimental dataset of *n* = 288 depolymerised images, using pure background as ground truth.

| **Metric** | **Value** |
| --- | --- |
| Brier Score | 6.2e-5 |
| BCE | 3.7e-4 |
| Weighted BCE (9:1) | 7.4e-5 |

The model performs even better on the depolymerised data, with the relevant losses multiple orders of magnitude lower than in the control case. Note that we do not include the AUC metrics in the depolymerised case, as the ground truth only covers a single class.

### 3.4 Tracking microtubule recovery

Using our trained model, we track the behaviour of the MT network in a single root hair cell over the experimental time course of MT depolyermsiation after Oryzalin treatment and its subsequent recovery after the Oryzalin drug has been washed out (Fig 7).

**Figure 7:**
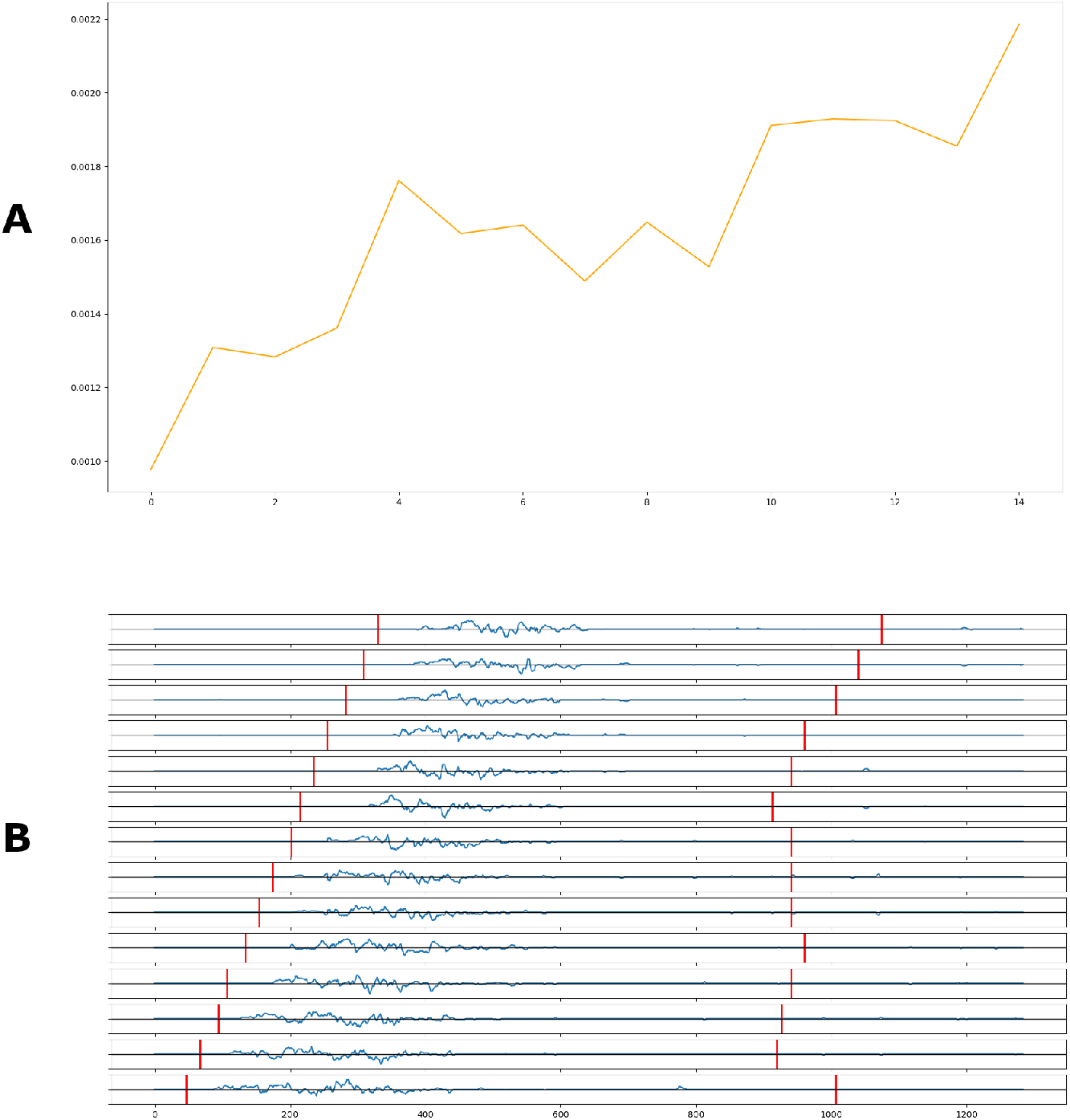
The evolution of the MT network as it recovers from effects of Oryzalin treatment. **A**, The total proportion of 3D images that are classified as MTs, plotted against time. **B**, The change in the intensity profiles of MTs between 3D images at consecutive time-steps. Time increases from top to bottom with the top image showing the difference between the 0th and 1st timesteps, and the last between the 13th and 14th timesteps. The x-axis corresponds to the long axis of the image, which correlates to the long axis of the root hair cell and is shown in pixels. The vertical red bars correspond to the tip of the cell (left) and the boundary of the nucleus (right).

For this example, we observe that the network recovers partially as shown by the incresee in the proportion of pixels registered as MTs increasing from a low of 0.1% to 0.2% (Fig 7A).

The MT network repolymerises closer to the tip than the nucleus (Fig 7B), and the polymerisation region traces the tip during growth. However, within this region there is no specific pattern to the repolymerisation.

## 4 Discussion and Conclusions

In this work, we generated realistic synthetic data to train a machine learning segmentation model, in order to quantify the depolymerisation and repolymerisation of microtubules in an *Arabidopsis* root hair cell from experimental images. Our novel pipeline to pre-process training images output from a computational microtubule model mimicked several biological processes such as microscopy optical effects, fluorophore distribution and MTs out of the plane. The generated synthetic images bore a close resemblance to experimental images at different stages of depolymerisation (Fig. 5), visually demonstrating that the pipeline could closely simulate experimental images. When this data was used for training, the model performed well on experimental images (Fig. 6). In particular, it was able to avoid segmenting the diffuse residue left behind by depolymerised MTs.

We validated the final model on unseen, experimental data. We considered the performance on control images (i.e. not exposed to OZ), where we used the outputs of Ilastik as a ground truth and also on completely depolymerised images, where the ground truth is solely background. We did not validate the model on experimental data in the intermediate stages of depolymerisation and repolymerisation, as we did not have access to a ground truth against which to compare. Using our model, we then ran inference on a single 3D time-series of a MT network repolymerising after Oryzalin was removed. The resulting intensity profiles suggests that the MT network repolymerises initially between the tip and nucleus.

Here we have focused on segmenting and analysing one set of experimental microtubule data. Future work could include generalising and testing this method on data collected in different microscopy conditions. While here we focus on microtubules, plant cells also contain another fibre network of actin. Future work could extend this segmentation methodology to actin. Preliminary testing, not presented here, suggests the model has low accuracy when used to segment actin data. The challenges may be similar to that of generalising to other microscope settings, as the fluorescent marker gives a different character of signal in the images of actin fibres. Alternatively, different training data may be required as the actin network behaviours are different to microtubules. For example, they can nucleate in different places. They also develop different structures to microtubules within the same cell which suggests different fibre behaviours or properties. Finally, it would also be interesting to extend to other plant cell types, focusing initially on other long thin cells such as trichomes or pollen tubes, and then in different organisms. Together, this future work would lead towards generating a single model which could segment cytoskeleton data of different quality and in different cell types

We have demonstrated that a machine learning tool trained on synthetic data generated from simulation outputs can segment experimental microtubule data in Arabidopsis cells. This opens up the potential for utilising simulation outputs for generating synthetic training data to develop next-generation image segmentation pipelines for fibre data, particularly in cases where experimental data is limited.

## Acknowledgments

K.E., T.A.S and H.J. acknowledge funding support from Gatsby through GAT3731/PR4. K.E. acknowledges support through the Cambridge Mathematics Placements (CMP) programme, and funded in part by donations made by corporate partners and individual donors to the CMP Supporters Club. We thank all members from the H.J. group for helpful discussions on this project particularly Nadia Radzam and Siyu Miao.

## Notes

https://gitlab.developers.cam.ac.uk/slcu/teamhj/publications/elangovan_etal_2025

### Competing Interest Statement

The authors have declared no competing interest.

